## Supplementary figures for "Post-replicative pairing of sister *ter* regions in *Escherichia coli* involves multiple activities of MatP"

### Supplementary information

Crozat et al.

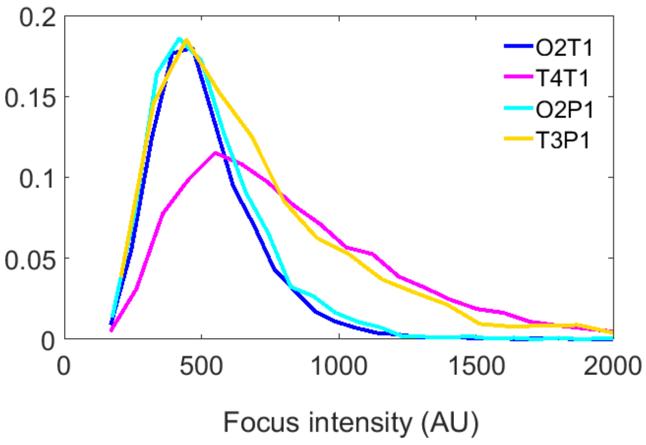


**Supplementary Figure 1: Intensities of foci labeled with ParS-P1 and ParS-pMT1**

Distribution of intensity for Ori2 and Ter3 or Ter4 foci labeled with ParS-P1 (O2P1 and T3P1) or ParS-pMT1 (O2T1 and T4T1). Note that ParB-P1 is fused with GFP, while ParB-pMT1 is fused with yGFP, which could result in a slight difference of intensity for the same number of GFP. Source data are provided as a Source Data file.

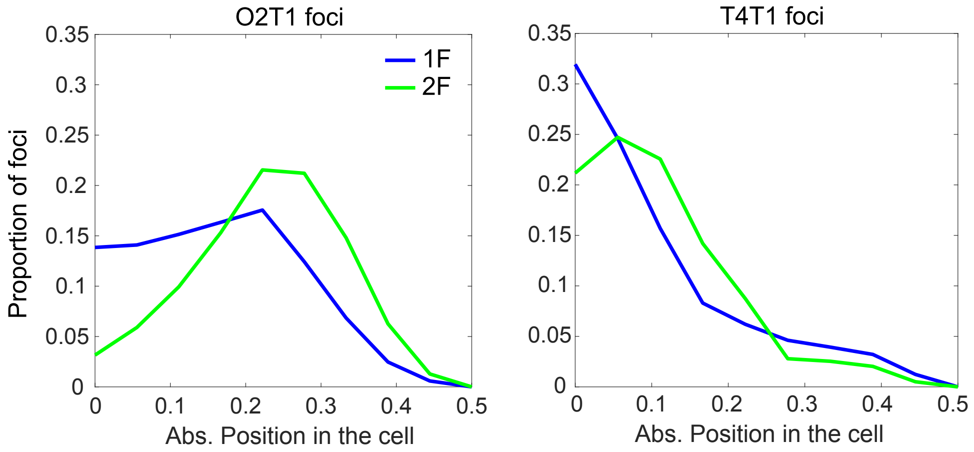


**Supplementary Figure 2: Cell localisation of Ori2 and Ter4 foci**

Localisation of Ter4 foci in a wild-type strain as a function of foci number and their intensity. Position in the cell is given from mid-cell (0) to the cell pole (0.5).

Note that only the foci that have a trajectory that has been validated by the script are plotted here, leading to an over-representation of 1F cells. 1F: 1 focus, 2F: 2 foci per cell. Source data are provided as a Source Data file.

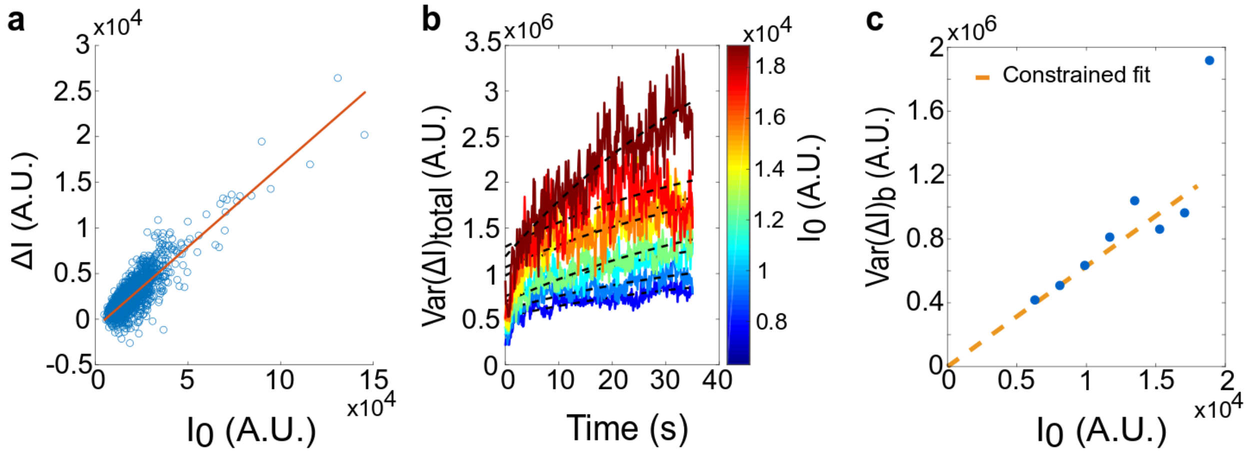


**Supplementary Figure 3: Calibration of intensity/copy number ratio**

**a** Background intensity, bleaching probability and time constant are estimated from the total intensity loss as a function of initial intensity. The estimated background intensity is subtracted before further processing. **b** Estimation of the bleaching contribution to variance over time. Tracks are binned by initial intensity, and variance in the intensity drop is computed per bin and per timepoint. Then, the variance as a function of time is fit with two terms, accounting for the bleaching and the short noise contributions to the intensity drop variance (see supplementary text for calculations). If variance was not dominated by bleaching, we would expect it to decay over time proportionally to intensity. On the graph, the bleaching term is roughly the difference between the end and the beginning of the black dashed curves. **c** The calibration parameter is estimated through a linear fit with intercept constrained to 0. The calibration parameter is obtained by dividing the slope of the linear fit by p(1-p), where p is the bleaching probability estimated in panel a. Source data are provided as a Source Data file.

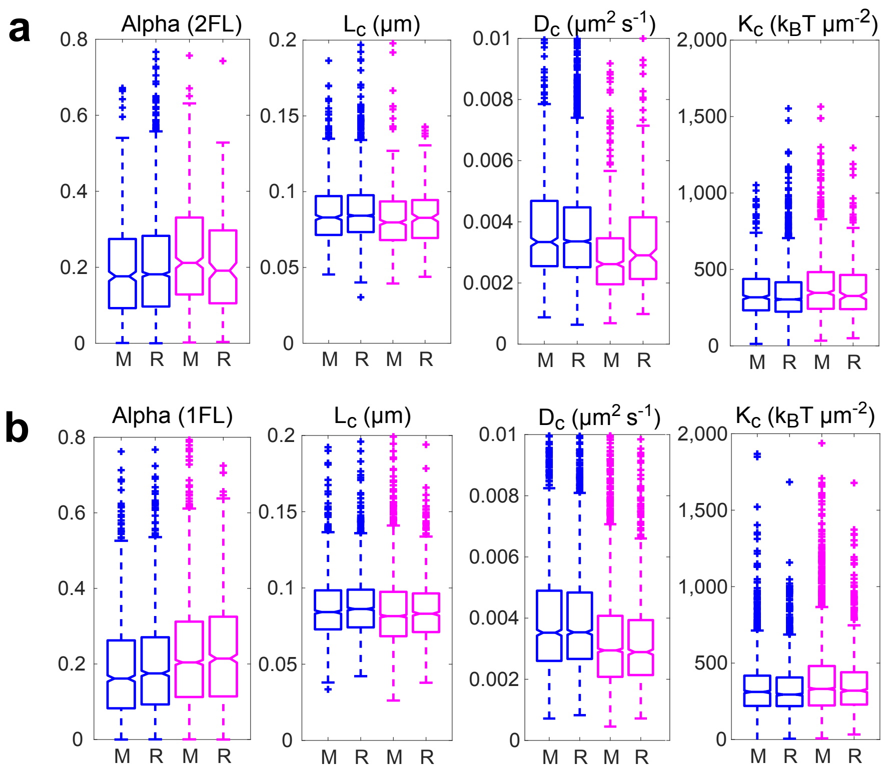


**Supplementary Figure 4: Mobility of 2FL in Ori2 and Ter4**

The four parameters α, $L_c$, $D_c$ and $K_c$ were calculated and are plotted as in Fig. 2D for Ori2 locus (blue) and Ter4 (pink) in wt background. **a** Results for 2FL. Ori2, n=639 for M, n=2716 for R; Ter4, n=588 for M, n=241 for R. **b** For an easier comparison, results are also shown for 1FL (same as in Fig. 2d). Ori2, n=1411 for M, n=1889 for R; Ter4, n=2744 for M, n=1064 for R. Box plots show the median of the distribution, the 25th and 75th percentiles of the samples (respectively the bottom and top of the box), the lowest and top values within a range of 1.5 times the interquartile range (dotted lines), and outliers of these as crosses. The notches display the variability of the median; non-overlapping notches between samples indicate different value of the medians at a 5% significance level. Source data are provided as a Source Data file.

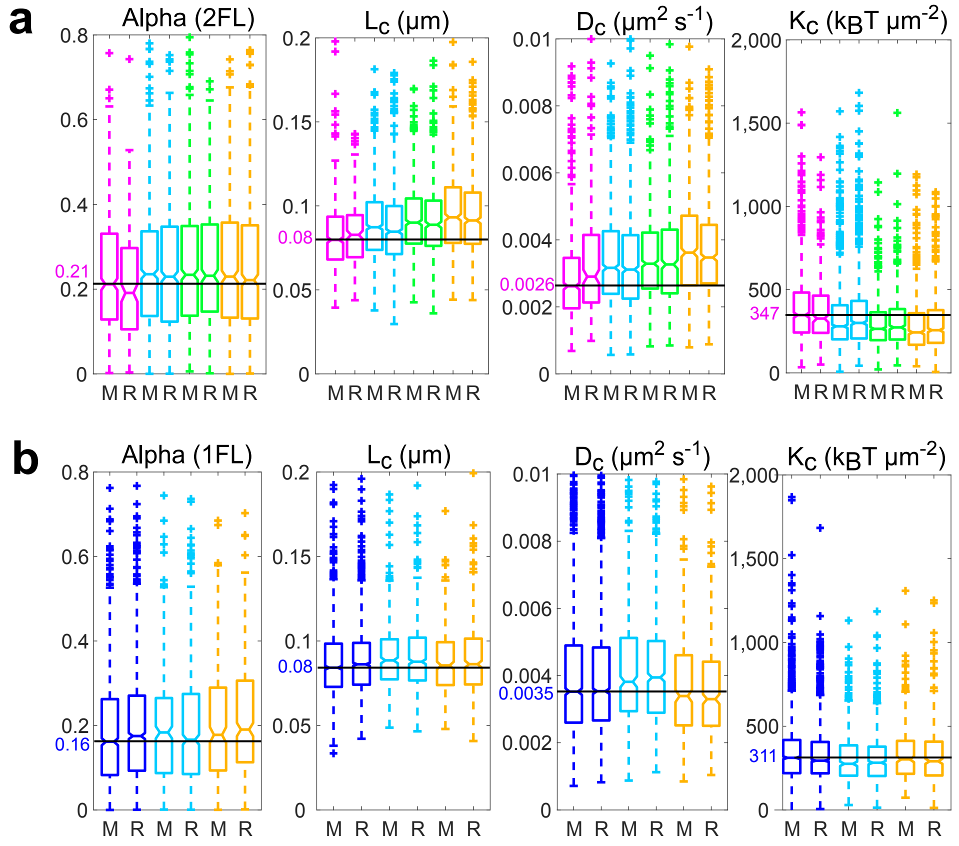


**Supplementary Figure 5: Mobility of FL foci in mutant strains**

The four parameters α, $L_c$, $D_c$ and $K_c$ were calculated and are plotted as in Fig. 2d. **a** Results for 2FL for Ter4 locus in wt (pink, n=588 for M, n=241 for R), Δ*matP* (light blue, n=1014 for M, n=979 for R), *matPΔ20* (green, n=813 for M, n=551 for R) and Δ*zapB* (orange, n=686 for M, n=733) backgrounds. **b** Ori2 locus in wt (dark blue, n=1411 for M, n=1889 for R), Δ*matP* (light blue, n=461 for M, n=766 for R) and Δ*zapB* (orange, n=328 for M, n=426 for R) backgrounds. Results here are shown for 1FL only, the number of 1FH being too low to give relevant results (Supplementary Table 1). To help comparing the values, a line has been drawn at the value for the foci at midcell in the wt background, the value indicated in pink (**a**) or blue (**b**), and the outliners at high values have been cut off. Box plots show the median of the distribution, the 25th and 75th percentiles of the samples (respectively the bottom and top of the box), the lowest and top values within a range of 1.5 times the interquartile range (dotted lines), and outliers of these as crosses. The notches display the variability of the median; non-overlapping notches between samples indicate different value of the medians at a 5% significance level. Source data are provided as a Source Data file.

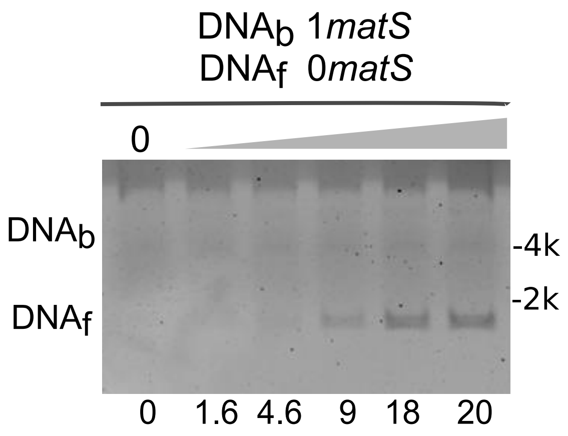


**Supplementary Figure 6: Pull-down experiment**

The experiment is done as in Fig. 4. Here $DNA_b$ contains one *matS*, while $DNA_f$ contains none. The percentage of DNA recovered is indicated at the bottom of the gel. The concentration of MatP is the same as in Fig. 5. Quantitation can vary from gel to gel but the trend is the same within 3 independent experiments. Molecular weight markers are indicated on the right of the gel (kbp).

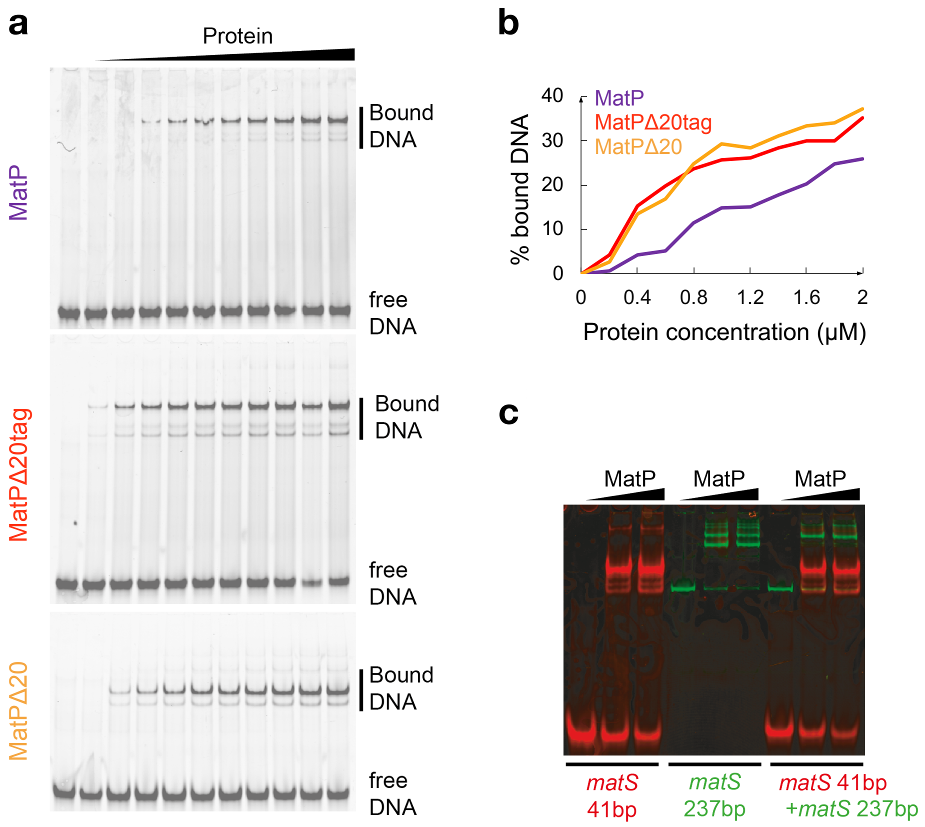


**Supplementary Figure 7: MatP interaction with DNA analysed by EMSA**

**a** EMSA experiment showing the interaction between an increasing concentration of protein (0.2, 0.4, 0.6, 0.8, 1, 1.2, 1.4, 1.6, 1.8, and 2μM) and a 41bp DNA fragment containing *matS*. Top: purified MatP protein. Middle: purified tagged MatPΔ20 protein. Bottom: purified untagged MatPΔ20 protein. **b** Percentage of bound DNA was estimated (ImageJ) and plotted as a function of the protein concentration. Source data are provided as a Source Data file. **c** EMSA experiment showing the interaction between MatP (0, 3 and 6 μM) and a 41bp, Cy3-labeled DNA molecule containing *matS* or a 237bp, Cy5-labeled DNA molecule containing *matS*. Note that a DNA-protein complex made of two DNA molecules of different length bridged together by MatP would appear yellow on this gel. These experiments were repeated three times.

**a**

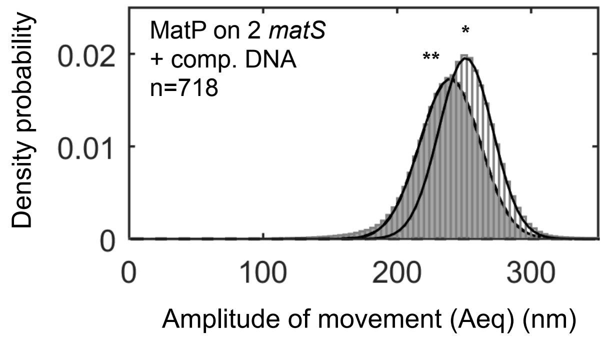


**b**

| DNA construct | Unlooped lifetime ($\tau_{fast}$, $\tau_{slow}$)(s) | Unlooped population | Number of events | Looped lifetime ($\tau_{fast}$, $\tau_{slow}$)(s) | Looped population | Number of events |
|---|---|---|---|---|---|---|
| 1 *matS* | 16.5 ±7.0<br>147.9±42.9 | 0.44<br>0.56 | 719 | 10.0 ±1.2<br>72.0±22.5 | 0.42<br>0.58 | 488 |
| 2 *matS* | 15.3 ±0.6<br>104.2±5.6 | 0.53<br>0.47 | 1,420 | 10.3 ±0.4<br>160.7±31.1 | 0.77<br>0.23 | 1,198 |

**c**

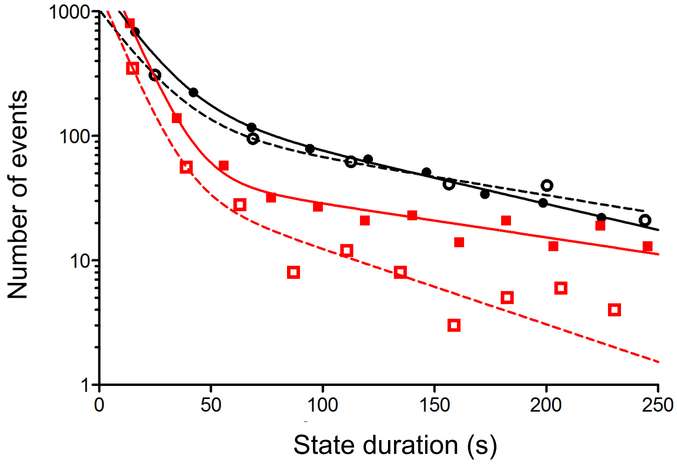


**Supplementary Figure 8: TPM with competitor DNA and kinetics analysis**

**a** Probability distributions of Aeq, before protein injection (light grey histogram), or during the 5min following MatP injection (dark grey histogram). The DNA contained 2 *matS*, 2.5µM of competitor DNA (dsDNA, 40bp, non-specific) was added in the reaction, and the number of tracks is 718. *, population centered on (mean +/- SD) : 251 +/-20nm; **, population centered on (mean +/- SD): 238 +/- 24nm. **b** Table of the Gaussian fit results of TPM time traces. **c** Histogram of the durations of unlooped (Black) and looped (Red) for DNA containing one matS (hollow symbols) or 2 matS (full symbols) in presence of MatP. Source data are provided as a Source Data file.

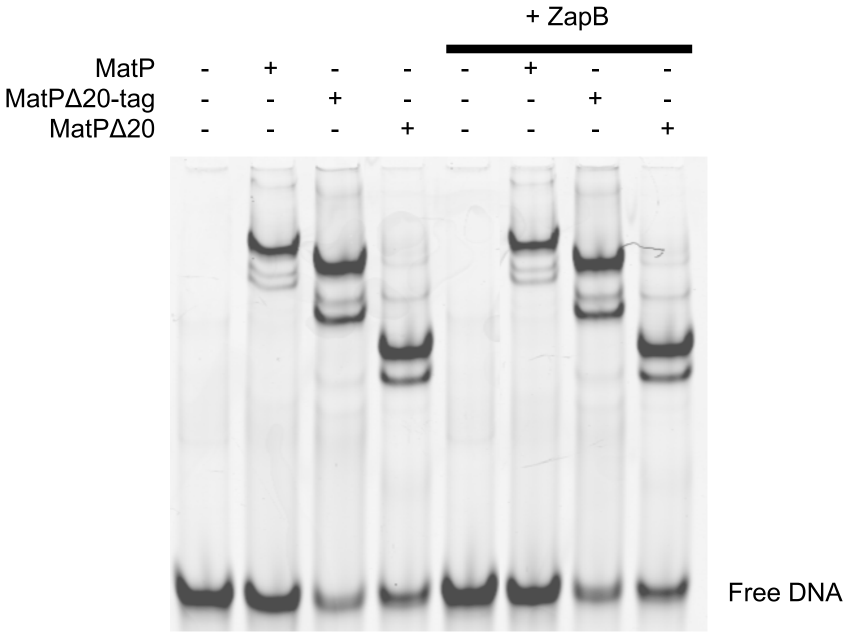


**Supplementary Figure 9: ZapB does not modify MatP/DNA interaction**

EMSA experiment showing the interaction between a 41bp DNA fragment containing *matS* and 3μM of MatP, his-tagged MatPΔ20, or untagged MatPΔ20. The experiment was done in absence of ZapB (first four lanes, also shown in Fig. 7) or in presence of 4μM of purified ZapB (last four lanes). This experiment was repeated twice, giving the same result.

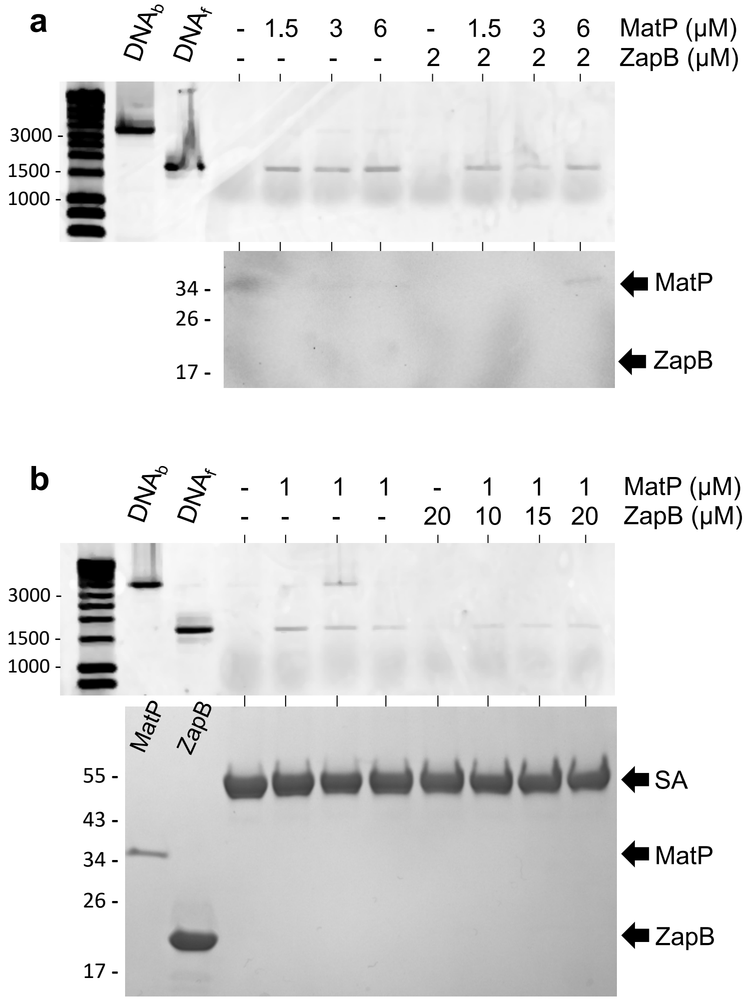


**Supplementary Figure 10: MatP/matS pull-down experiments in presence of ZapB**

This experiment is performed as explained in Fig. 4. $DNA_b$ is associated to a magnetic bead whereas $DNA_f$ is not. Both contains 2 *matS* sites. **a** The top gel (1% agarose gel electrophoresis) shows the pull-down of $DNA_f$ when different concentration of MatP are added (1.5; 3 or 6μM). The same pull-down is observed when 2μM of ZapB are also added in the reaction. The bottom gel corresponds to the same pull-down experiment but analysed by SDS-PAGE, Western blot and anti-His immuno-detection. No ZapB is detected. **b** The top gel (1% agarose gel electrophoresis) shows the pull-down of $DNA_f$ when 1μM of MatP is added. The same pull-down is observed when large amount of ZapB are added (10μM, 20μM or 30μM). The bottom gel corresponds to the same pull-down experiment but analysed by SDS-PAGE, and stained with Coomassie blue. No ZapB is detected. The first two lanes show a sample of purified MatP and ZapB. SA, streptavidin. Size markers are indicated on the left of each gel, in bp for agarose gels and kDa for protein gels. Both experiments were repeated twice and gave the same results.

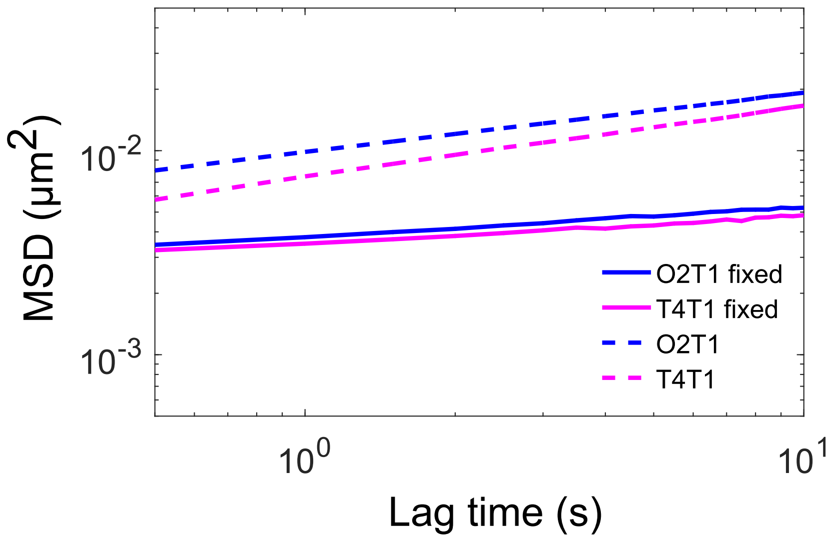


**Supplementary Figure 11: Mobility of foci in fixed cells**

To assess the limitation of our detection system, a control was performed with fixed cells. Ori2 (blue curves) and Ter4 (pink curves) foci were followed as usual, and their MSD was calculated. The dotted lines show the medians of MSDs of foci for the non-fixed cells used on the same day. Source data are provided as a Source Data file.

**Supplementary Table 1: List of primers used in this study**

| Name | F/R | 5'-3' Sequence | Matrix |
|---|---|---|---|
| **OEB3** | Rv | b-GGTCGTGAAGCGATTCACA | pBR322 - 2matS |
| **OEB4** | Fw | GTAGATAACTACGATACGGG | pBR322 - 2matS |
| **OEB5** | Fw | GCCACAGTCGATGAATCCA | pBR322 - 2matS |
| **OEB6** | Rv | b-CCCGTATCGTAGTTATCTAC | pBR322 - 2matS |
| **OEB7** | Rv | GAATTCTTGAAGACGAAAGGG | pBR322 - 2matS |
| **OEB8** | Fw | b-TTGAGAGCCTTCAACCCAGT | pBR322 - 2matS |
| **OEB9** | Fw | GGGTTATTGTCTCATGAGCG | pBR322 - 2matS |
| **OEB10** | Rv | TCCAGCGAAAGCGGTCC | pBR322 - 2matS |
| **OEB11** | Rv | GTGAATCCGTTAGCGAGGT | pBR322 - 2matS |
| **OEB12** | Fw | TTACCGGATACCTGTCCGC | pBR322 - 2matS |
| **matS41F** | Fw | CY3-AAGTACGGTAAAAGGTGACAGTGTCACTTTCATTGTTGGTA | N/A |
| **matS41R** | Rv | TACCAACAATGAAAGTGACACTGTCACCTTTTACCGTACTT | N/A |
| **matSF** | Fw | CY5-GTAGTGCCGGGAGAAAGCAG | pEL3 |
| **matSR** | Rv | GCCTTCTTGACGAGTTCTTC | pEL3 |
| **F2060** | Fw | b-CTGCAATGATACCGCGAGAC | pBR328/pTOC7 |
| **R2060** | Rv | dig-TGACTTCCGCGTTTCCAGAC | pBR328/pTOC7 |
| **F1201** | Fw | b-CTGGTGAGTACTCAACCAAG | pTOC6 |
| **R1201** | Rv | dig-CTACAATCCATGCCAACC | pTOC6 |

b- biotinylated oligo
CY3- CY3 labeled oligo
CY5- CY5 labeled oligo
Dig- digoxigenine labeled oligo
